# Determinants of base-pair substitution patterns revealed by whole-genome sequencing of DNA mismatch repair defective *Escherichia coli*

**DOI:** 10.1101/346874

**Authors:** Patricia L. Foster, Brittany A. Niccum, Ellen Popodi, Jesse P. Townes, Heewook Lee, Wazim MohammedIsmail, Haixu Tang

**Author notes:** Current address: Computational Biology Department, School of Computer Science, Carnegie Mellon University, Pittsburgh, PA, USA, 15213. **Corresponding author:** Patricia L Foster, Ph.D., Department of Biology, Indiana University, 1001 E. Third St, Bloomington, IN 47405 USA.

## Abstract

Mismatch repair (MMR) is a major contributor to replication fidelity, but its impact varies with sequence context and the nature of the mismatch. Mutation accumulation experiments followed by whole-genome sequencing of MMR-defective *E. coli* strains yielded ≈30,000 base-pair substitutions, revealing mutational patterns across the entire chromosome. The base-pair substitution spectrum was dominated by A:T > G:C transitions, which occurred predominantly at the center base of 5′N**A**C3′+5′G**T**N3′ triplets. Surprisingly, growth on minimal medium or at low temperature attenuated these mutations. Mononucleotide runs were also hotspots for base-pair substitutions, and the rate at which these occurred increased with run length. Comparison with ≈2000 base-pair substitutions accumulated in MMR-proficient strains revealed that both kinds of hotspots appeared in the wild-type spectrum and so are likely to be sites of frequent replication errors. In MMR-defective strains transitions were strand biased, occurring twice as often when A and C rather than T and G were on the lagging-strand template. Loss of nucleotide diphosphate kinase increases the cellular concentration of dCTP, which resulted in increased rates of mutations due to misinsertion of C opposite A and T. In an *mmr ndk* double mutant strain, these mutations were more frequent when the template A and T were on the leading strand, suggesting that lagging-strand synthesis was more error-prone or less well corrected by proofreading than was leading strand synthesis.

## INTRODUCTION

In 1961 Seymour Benzer published a paper entitled “On the topography of the genetic fine structure” (Benzer 1961). Using recombinational mapping of 1,612 independent spontaneous mutations in the *rII* gene of bacteriophage T4, Benzer revealed that not all sites in the gene were equally mutable; over 500 frameshift mutations were recovered at one site whereas about 200 sites had only 1 or 2 frameshift mutations. Subsequent sequencing showed that Benzer’s hotspot was a run of 6 As, as were two other hotspots in the gene (Pribnow *et al.* 1981), and such iterated sequences were readily found to be hotspots for small insertions and deletions (indels) in other model systems (Streisinger *et al.* 1966; Farabaugh *et al.* 1978). The most likely reason for this phenomenon is that iterated sequences are “slippery”, *i.e.* are places where DNA strands can misalign, causing small indels during replication (Kunkel 2004).

The occurrence of hotspots is not restricted to indels, nor to spontaneous mutations. Studies of mutagenesis over the years have revealed that, both on a small- and on a large-scale, different segments of DNA vary widely in mutability. Yet the understanding of what determines these differences, particularly in the case of point mutations, are limited to a few special cases, such as methylated bases (Coulondre *et al.* 1978; Lee *et al.* 2012), quasipalindromic sequences (de Boer and Ripley 1984; Viswanathan *et al.* 2000), and the repeat sequences mentioned above. Even among sites within the same gene with the same local sequence context, base-pair substitution (BPS) rates can vary by more than an order of magnitude (Garibyan *et al.* 2003). And, at least in mismatch repair (MMR) defective lines, BPS rates vary 2 to 3-fold across bacterial genomes in wave like patterns symmetrical about the origin (Foster *et al.* 2013; Dettman *et al.* 2016; Dillon *et al.* 2017). Thus, there is much yet to learn about what determines the mutability of any given DNA site.

Another lesson learned from Benzer’s classic paper is the importance of large numbers when studying rare events such as mutations. Investigating mutational processes with mutation accumulation (MA) protocols that exert minimal selective pressure has the advantage of allowing mutations to accumulate in an unbiased manner. Coupling the MA protocol to whole-genome sequencing (WGS) eliminates the possibility that peculiarities of particular DNA segments can bias the overall results, and also allows mutation rates across the genome to be evaluated. However, the laboriousness of the MA procedure limits the numbers of mutations that can be analyzed. To overcome this limitation, we and others have used mutator strains of model microorganisms.

DNA replication in E. coli is performed by DNA polymerase III holoenzyme, a multisubunit machine with high processivity and accuracy. The fidelity of replication, which, in *E. coli*, is about 1 mistake in 1000 generations (Lee *et al.* 2012), is mainly due to three factors: the intrinsic base-pairing fidelity of the DNA polymerase, error-correction by the exonuclease activity of the proofreader, and correction of mismatches by the mismatch-repair system (MMR). In *E. coli*, mismatch repair is accomplished by four major enzymes. MutS surveys the DNA after replication, finds mismatches and binds to them. MutS then recruits MutL, and together they find a nearby GATC site, which, in *E. coli*, is methylated at the A on the template, “old,” DNA strand. MutS and MutL then recruit MutH and activated it to nick the DNA on the unmethylated, “new,” DNA strand. The UvrD helicase, together with one of four exonucleases, removes the nicked strand past the mismatch. Pol III holoenzyme then assembles and synthesizes a new strand that, most likely, has the correct sequence. MMR is highly conserved between bacteria and eukaryotes and, in general, improves replication fidelity by 100-fold or more (reviewed in (Marinus 2012; Ganai and Johansson 2016).

In the studies reported here, we have combined the results of 10 separate MA/WGS experiments with MMR-defective *E. coli* strains to obtain a collection of 30,061 BPS, giving a data base with strong statistical power. In addition, combining 8 MA/WGS experiments with MMR-proficient strains yielded 1,933 BPS with which to make some statistically significant comparisons. We focused on BPS because, in our MA experiments, they are the most prominent class of spontaneous mutations, occurring at 10-times the rate as small insertions and deletions (indels) in wild-type cells, and 5-times the rate of indels in MMR defective cells (Lee *et al.* 2012; Foster *et al.* 2015). The results give insights into the determinants of spontaneous rates of BPSs in different genetic backgrounds, under different growth conditions, and at different DNA sites.

## MATERIALS AND METHODS

### Bacterial strains and media

The bacterial strains used and the methods of their construction are given in Supplemental Table S1. The oligonucleotides used to perform and confirm genetic manipulations are given in Supplemental Table S2. Further details are given in Supplemental Materials and Methods. For analysis of wild-type mutation rates and spectra, the results from MA experiments with the following strains were combined: PFM2, wild-type, 3K and 6K (Lee *et al.* 2012); PFM35, *uvrA*; PFM40, *alkA tagA*; PFM88, *ada ogt*; PFM91, *nfi*; PFM101, *umuDC dinB*; PFM133, *umuDC dinB polB*; the mutation rates and spectra of all of these strains were equivalent (Foster *et al.* 2015).

### Mutation accumulation procedure

The MA procedure was as described (Lee *et al.* 2012; Foster *et al.* 2015). The MA lines originated from two or more founder colonies and were propagated through single-colony bottlenecks as described (Lee *et al.* 2012). Further details are given in Supplemental Materials and Methods.

### Estimation of generations

The number of generations that each MA line experienced was estimated from colony size as previous described (Lee *et al.* 2012). Further details are given in Supplemental Materials and Methods.

### Estimation of mutation rates

For each MA experiment the mutation rate was estimated by dividing the total number of mutations accumulated by all the MA lines by the total number of generations that were undergone. This value for mutations per generation was then divided by the appropriate number of sites (A:T sites, G:C sites, etc) to give the conditional mutation rate. The individual mutation rates for each line were used in statistical analyses (see below).

Estimation of mutation rates from fluctuation tests is described in Supplemental Materials and Methods.

### Determination of DNA strand bias

In *E. coli*, replication is bidirectional starting at the origin and proceeding through right and left halves of the chromosome, called replichores. The reported DNA strand in sequenced genomes is the 5′ to 3′ reference, or “top”, strand, which is the lagging-strand template (LGST) on the right replichore, and the leading strand template (LDST) on the left replichore. For example, the reference strand triplet 5′T**A**C3′ (with the center base the mutational target) on the right replichore has the target purine on the LGST and the target pyrimidine on the LDST, whereas the same triplet on the left replichore has the opposite orientation. Therefore, 5′T**A**C3′ on the right replichore plus 5′G**T**A3′ on the left replichore are both instances of that triplet with the target purine on the LGST and the target pyrimidine on the LDST. Likewise 5′T**A**C3′ on the left replichore plus 5′G**T**A3′ on the right replichore are both instances of that triplet with the target purine on the LDST and the target pyrimidine on the LGST. To calculate the strand biases of BPSs, the number of BPSs at each LGST and LDST triplet was divided by the number these triplets in the genome.

### Genomic DNA preparation, library construction, sequencing, and SNP calling

Genomic DNA was purified using the PureLink Genomic DNA purification kit (Invitrogen Corp.). Libraries were constructed by the Indiana Univ. Center for Genomics and Bioinformatics (IUCGB). Sequencing of most experiments was performed by Univ. of New Hampshire Hubbard Center for Genome Studies using the Illumina HiSeq 2500 platform. For the experiment at low temperature (see below), sequencing was done by the IUCGB using the Illumina NextSeq platform. Procedures for SNP and indel calling were as described (Lee *et al.* 2012). NCBI Reference Sequence, NC_000913.2 (*E. coli* K12 strain MG1655) was used as the reference genome sequence. Further details are given in Supplemental Materials and Methods.

### Mutation annotation

Variants were annotated using custom scripts. Protein coding gene coordinates were obtained from the GenBank page of reference sequence, NC_000913.2. BPSs in coding sequences were determined to be synonymous or nonsynonymous based on the genetic code. Nonsynonymous BPSs were designated conservative or nonconservative based on the Blosum62 matrix (Henikoff and Henikoff 1994) with a value ≥0 considered conservative.

### Statistical analyses

Values and 95% CL for ratios between variables were calculated as in Rice, 1995 (Rice 1995). Other statistical tests were performed as in Zar, 1984 (Zar 1984). To compare results among MA lines and among experiments, the numbers of mutations had to be normalized by the number of generations. To do these comparisons, we treated the mutations per generation as weighted variables and used the following formulae, adapted from Mandansky, 2010 (Mandansky 2010), to calculate the variance (Var).

For each experiment:

m = mutations per MA line; g = generations per MA line; mutation rate of a MA line = r = m/g Mean overall mutation rate = R = M/G where M=Σm and G=Σg over all MA lines.

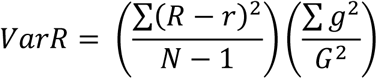

where N = number of MA lines being considered. Then

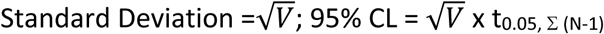

To combine the results from multiple experiments, the same calculations were used except (N1) was replaced with Σ(N-1), i.e. (N-1) summed over all experiments.

### Data availability

Strains are available upon request. File S1 contains supplemental Materials and Methods. File S2 contains supplemental Tables, which include strain genotypes and methods of construction, oligonucleotide sequences, and detailed data from each experiment. File S3 contains supplemental figures that are referenced in the text. The sequences and SNPs reported in this paper have been deposited in the National Center for Biotechnology Information Sequence Read Archive (BioProject accession no. TBA) and in the IUScholarWorks Repository (URL: TBA).

## RESULTS

### Transitions at A:T basepairs dominate the base-pair substitution (BPS) spectrum in mismatch-repair (MMR) defective strains

In a previous paper we reported that in a *mutL* defective *E. coli* strain, the spectrum of BPSs is dominated by A:T to G:C transitions (Lee *et al.* 2012). Because this result was contrary to some previous studies reporting that loss of MMR increases the rates of A:T and G:C transitions equally (Choy and Fowler 1985; Cupples and Miller 1988), we repeated MA experiments with additional strains deleted for each of the genes that encode the proteins of the MMR system. As shown in Figure 1A&B, both the mutation rates and the mutational spectra of strains deleted for one, two, or all of the genes that encode the MutSLH complex were nearly the same. In contrast, although the mutational spectrum was the same, the mutation rate of a strain missing UvrD, the helicase that works with the MMR system, was about half that of the strains defective for MutSLH (Figure 1C&D). This result implies that another helicase, possibly Rep, an UvrD homolog, can perform this function.

**Figure 1.**
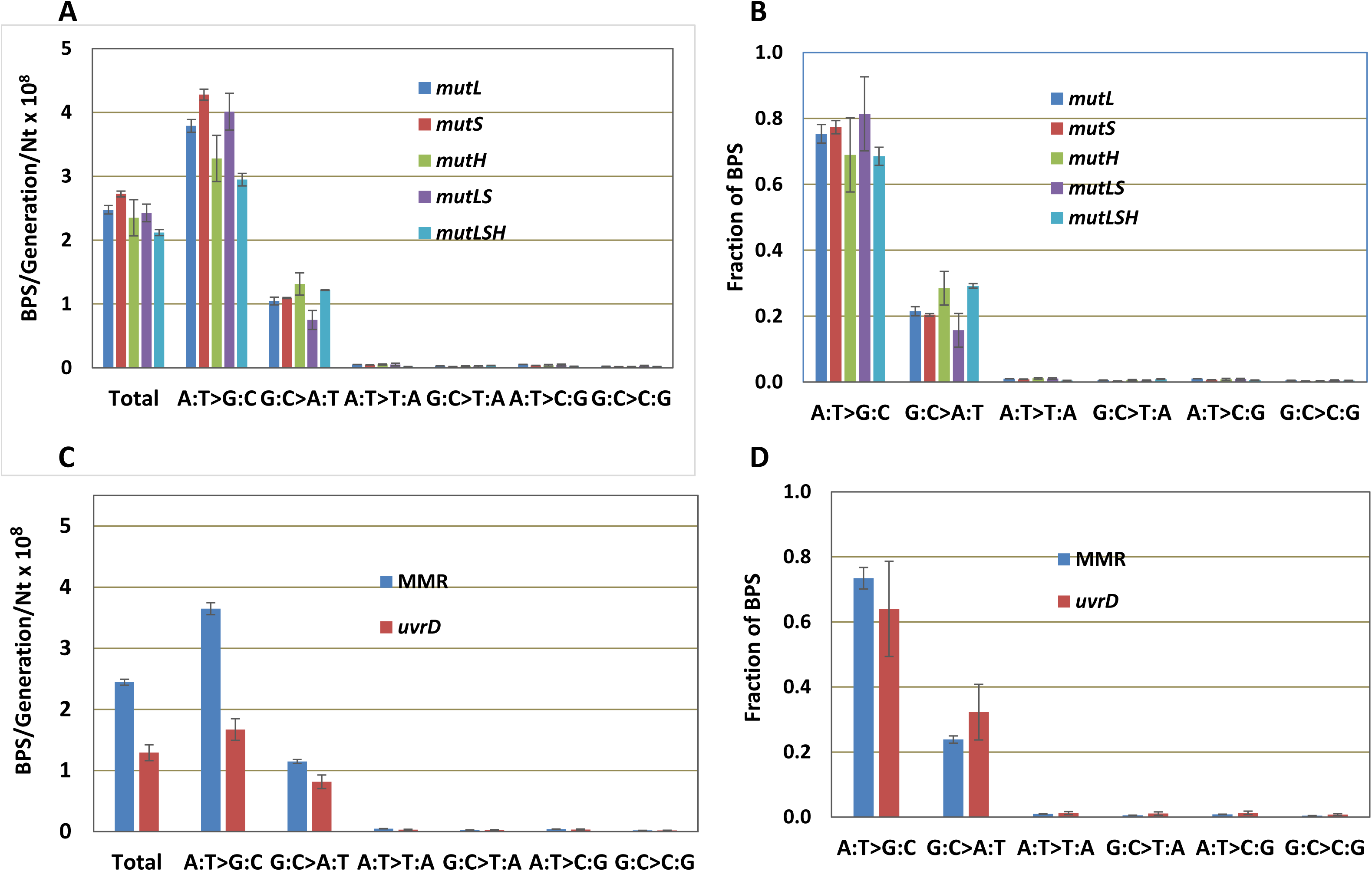
All MMR-defective strains have the same mutational spectrum but *uvrD* mutant strains have a lower mutation rate. A & B. The data for all MA experiments with strains with the same MMR defect have been combined (see Materials and Methods); the number of experiments with each are: *mutL*, three; *mutS*, three; *mutH*, one; *mutLS*, one; *mutLSH*, two. The results of each MA experiment are given in Supplemental Tables S3, S4, and S5. C&D. The data from the ten MA experiments with strains defective for *mutL*, *mutS*, and *mutH* are combined to give the results labeled MMR (see Materials and Methods). Bars represent the means and error bars are 95% CL for both rates and fractions. BPS, basepair substitution; Nt, nucleotide

Because of the uniformity of results with the MutSLH-defective strains, the data from the 10 independent experiments, consisting of 334 MA lines, have been combined in Figure 1C&D and the analyses to follow, and appear labeled as “MMR” (see Supplemental Tables S3, S4, and S5 for the results of each experiment). In all, these strains underwent almost 250,000 generations, accumulating 30,061 BPSs, resulting in a data set with great statistical power.

We investigated several hypotheses to explain the dominance of A:T transitions (Table 1). *E. coli* has two error-prone polymerases, DNA Pol IV and V, that could be responsible for these mutations, but deleting the genes that encode both polymerases had little effect on the BPS rate and no effect on the spectrum of the MMR defective strain. MutY is a glycosylase that removes As mispaired with Gs or 8-oxoGs as part of the pathway that protects cells from the mutagenic effects of oxidation. MutY also has activity against As mispaired with Cs, which creates A:T to G:C transitions if the A is the correct base (Kim *et al.* 2003); this second activity of MutY could explain the dominance of A:T transitions in the spectrum of the MMR-defective strains. However, the only major effect of loss of MutY in the *mutL* deletion strain was an increase in the rate of G:C to T:A mutations, as expected. Mfd is the factor that initiates transcription-coupled repair (TCR), a pathway that preferentially repairs damage to the transcribed DNA strand during active transcription (reviewed in (Ganesan *et al.* 2012). While TCR is performed by the proteins that participate in nucleotide excision repair (Selby and Sancar 1993), some reports have also implicated MutS and/or MutL in TCR (Li and Bockrathi 1995; Mellion and Champe 1996). We hypothesized that in absence of MMR, TCR was repairing G:C to A:T mutations, or (less likely), creating A:T to G:C mutations, resulting in the observed mutational bias. But, deleting Mfd had little effect on the BPS rate or spectrum of the MMRdefective strain.

**Table 1A:** Conditional BPS rates in strains grown on different media and at different temperatures

| Strain | Genotype | Medium | Total | Mutations/Generation/Nt x 10<sup>8</sup> ± 95% CL |  |  |  |  |  |
| --- | --- | --- | --- | --- | --- | --- | --- | --- | --- |
|  |  |  |  | A:T>G:C | G:C>A:T | A:T>T:A | G:C>T:A | A:T>C:G | G:C>C:G |
| *MMR | *mmr* | LB | 2.45 ± 0.05 | 3.65 ± 0.10 | 1.15 ± 0.03 | 0.05 ± 0.002 | 0.03 ± 0.001 | 0.04 ± 0.002 | 0.02 ± 0.001 |
| PFM118 | *mutL umuDC dinB* | LB | 1.98 ± 0.22 | 2.89 ± 0.33 | 0.99 ± 0.16 | 0.04 ± 0.03 | 0.02 ± 0.02 | 0.03 ± 0.02 | 0.02 ± 0.02 |
| PFM137 | *mutL mutY* | LB | 2.89 ± 0.17 | 4.39 ± 0.32 | 0.97 ± 0.14 | 0.06 ± 0.03 | 0.36 ± 0.06 | 0.03 ± 0.02 | 0.02 ± 0.02 |
| PFM294 | *mutL mfd* | LB | 2.34 ± 0.21 | 3.82 ± 0.40 | 0.81 ± 0.09 | 0.04 ± 0.02 | 0.02 ± 0.01 | 0.04 ± 0.02 | 0.02 ± 0.01 |
| PFM368 | *mutS mfd* | LB | 3.02 ± 0.17 | 4.75 ± 0.30 | 1.21 ± 0.14 | 0.06 ± 0.02 | 0.02 ± 0.01 | 0.05 ± 0.02 | 0.02 ± 0.01 |
| PFM5 | *mutL* | Minimal | 1.10 ± 0.07 | 1.03 ± 0.09 | 1.11 ± 0.11 | 0.01 ± 0.01 | 0.02 ± 0.01 | 0.01 ± 0.01 | 0.02 ± 0.01 |
| PFM137 | *mutL mutY* | Minimal | 2.17 ± 0.17 | 1.26 ± 0.15 | 1.45 ± 0.19 | 0.02 ± 0.02 | 1.54 ± 0.19 | 0.01 ± 0.01 | 0.03 ± 0.02 |
| PFM144 | *mutL* | Buffered LB | 2.08 ± 0.20 | 2.82 ± 0.37 | 1.23 ± 0.13 | 0.04 ± 0.03 | 0.03 ± 0.02 | 0.04 ± 0.02 | 0.03 ± 0.02 |
| #PFM343 | *mutS* | LB | 2.75 ± 0.14 | 4.25 ± 0.25 | 1.15 ± 0.12 | 0.06 ± 0.03 | 0.03 ± 0.01 | 0.04 ± 0.01 | 0.02 ± 0.01 |
| #PFM343 | *mutS* | Diluted LB | 3.40 ± 0.17 | 4.29 ± 0.28 | 2.37 ± 0.19 | 0.06 ± 0.02 | 0.04 ± 0.02 | 0.04 ± 0.02 | 0.03 ± 0.02 |
| #PFM343 | *mutS* | Supplemented<br>Minimal | 2.32 ± 0.14 | 2.43 ± 0.17 | 2.08 ± 0.17 | 0.03 ± 0.01 | 0.05 ± 0.04 | 0.03 ± 0.01 | 0.02 ± 0.01 |
| #PFM343 | *mutS* | Minimal | 1.32 ± 0.08 | 1.27 ± 0.11 | 1.30 ± 0.11 | 0.02 ± 0.01 | 0.02 ± 0.01 | 0.005 ± 0.007 | 0.02 ± 0.01 |
| #PFM342 | *mutS* | †LB, 28°C | 1.09 ± 0.10 | 1.31 ± 0.13 | 0.85 ± 0.12 | 0.01 ± 0.01 | 0.01 ± 0.01 | 0.01 ± 0.01 | 0.01 ± 0.01 |

**Table 1B:** Fractions of BPSs strains grown on different media and at different temperatures

| Strain | Genotype | Medium | BPS Rate/Total Rate ± 95% CL | | | | | |
| --- | --- | --- | --- | --- | --- | --- | --- | --- |
|  |  |  | A:T>G:C | G:C>A:T | A:T>T:A | G:C>T:A | A:T>C:G | G:C>C:G |
| *MMR | *MMR | LB | 0.73 ± 0.23 | 0.24 ± 0.08 | 0.01 ± 0.005 | 0.005 ± 0.003 | 0.008 ± 0.004 | 0.004 ± 0.002 |
| PFM118 | *mutL umuDC dinB* | LB | 0.72 ± 0.16 | 0.26 ± 0.07 | 0.01 ± 0.01 | 0.01 ± 0.01 | 0.01 ± 0.01 | 0.01 ± 0.01 |
| PFM137 | *mutL mutY* | LB | 0.75 ± 0.09 | 0.17 ± 0.04 | 0.01 ± 0.01 | 0.06 ± 0.02 | 0.005 ± 0.005 | 0.003 ± 0.005 |
| PFM294 | *mutL mfd* | LB | 0.80 ± 0.14 | 0.18 ± 0.03 | 0.01 ± 0.01 | 0.004 ± 0.004 | 0.01 ± 0.01 | 0.003 ± 0.004 |
| PFM368 | *mutS mfd* | LB | 0.77 ± 0.08 | 0.20 ± 0.03 | 0.01 ± 0.004 | 0.004 ± 0.003 | 0.01 ± 0.004 | 0.003 ± 0.002 |
| PFM5 | *mutL* | Minimal | 0.46 ± 0.11 | 0.52 ± 0.13 | 0.004 ± 0.007 | 0.01 ± 0.01 | 0.003 ± 0.006 | 0.01 ± 0.01 |
| PFM137 | *mutL mutY* | Minimal | 0.29 ± 0.01 | 0.35 ± 0.02 | 0.01 ± 0.002 | 0.37 ± 0.02 | 0.001 ± 0.001 | 0.01 ± 0.002 |
| PFM144 | *mutL* | Buffered LB | 0.67 ± 0.16 | 0.30 ± 0.06 | 0.01 ± 0.01 | 0.01 ± 0.01 | 0.01 ± 0.01 | 0.01 ± 0.01 |
| #PFM343 | *mutS* | LB | 0.76 ± 0.06 | 0.21 ± 0.02 | 0.01 ± 0.005 | 0.01 ± 0.002 | 0.01 ± 0.003 | 0.004 ± 0.002 |
| #PFM343 | *mutS* | Diluted LB | 0.62 ± 0.08 | 0.35 ± 0.05 | 0.01 ± 0.004 | 0.01 ± 0.003 | 0.01 ± 0.003 | 0.004 ± 0.003 |
| #PFM343 | *mutS* | Supplemented<br>Minimal | 0.52 ± 0.09 | 0.46 ± 0.09 | 0.01 ± 0.01 | 0.01 ± 0.02 | 0.02 ± 0.02 | 0.004 ± 0.005 |
| #PFM343 | *mutS* | Minimal | 0.48 ± 0.10 | 0.50 ± 0.11 | 0.01 ± 0.01 | 0.01 ± 0.01 | 0.002 ± 0.006 | 0.01 ± 0.01 |
| #PFM342 | *mutS* | †LB, 28°C | 0.59 ± 0.08 | 0.39 ± 0.06 | 0.002 ± 0.003 | 0.005 ± 0.003 | 0.003 ± 0.003 | 0.006 ± 0.004 |

\*10 experiments with MMR-defective strains grown on LB have been combined (see Supplemental Tables S3 and S4).
#PFM342 and PFM343 were constructed in parallel had have the same phenotype (see Supplemental Tables S2, S3, and S4
†For all other experiments strains were grown at 37°C.

### Different growth media and temperatures produce different BPS spectra

Surprisingly, growing MMR-defective strains on glucose minimal medium completely eliminated the excess A:T transitions, resulting in nearly equal rates for both transitions (Table 1). One important difference between LB and minimal medium is that the latter is buffered at pH 7, whereas the pH of LB, although initially adjusted to 7, first falls and then rises as the cells grow (Luli and Strohl 1990; McFall and Newman 1996). To test whether these pH changes were responsible for the excess A:T to G:C mutations in LB, we repeated the MA experiment with the LB buffered to pH 7. However, the results were the same as when the cells were grown on unbuffered LB (Table 1).

Two additional interconnected factors could affect the mutational results obtained on the two media. First, the growth rate of cells on minimal medium, as measured by colony size, is about half of that on LB. A slower growth rate, reflecting slower replication, would give DNA polymerase more time to insert the correct base, and proofreader more time to remove an incorrect base. Second, the physiological state of the cells is greatly different in the two media: cells in LB are mainly metabolizing peptides and amino acids, whereas cells in our minimal medium are utilizing glucose and must synthesize all their amino acids. To evaluate these factors we first performed MA experiments using two additional media: LB medium diluted 1:5, which slowed the growth rate of the cells to about that on minimal medium, and minimal glucose medium supplemented with LB at a ratio of 10:1, which increased the growth rate of the cells to about that on LB (Supplemental Table S6). As shown in Table 1, the results were ambiguous. The almost 4-fold ratio of A:T to G:C transitions observed in cells grown on LB was reduced to 2-fold when the LB was diluted, whereas the 1:1 ratio of A:T to G:C transitions observed in cells grown on minimal medium became a 1.2-fold excess of AT transitions when the minimal medium was supplemented with LB.

To further distinguish between the effects of growth rate and medium, we performed a MA experiment using LB, but incubating the plates at 28°C, which resulted in the cells growing at about the same rate as they did on minimal glucose medium (Supplemental Table S6). The resulting mutation rate was 40% that of the rate at 37°C on LB, which was close to the rate of cells on minimal glucose medium. There were 70% fewer A:T transitions at low temperature than at 37°C on LB, but G:C transitions also dropped by 30%, so the ratio of A:T to G:C transitions, 1.5, was again an intermediate number. But since the decline in A:T transitions was more than twice the decline in G:C transitions, it would appear that the growth rate is the dominant factor determining the rate of A:T transitions.

Unexpectedly, when the LB was diluted, the mutation rate increased by about 25% relative to the rate on undiluted LB. This increase was due entirely to a 2-fold increase in the rate of G:C transitions, which could indicate that LB also has a suppressive effect on these mutations. Thus the domination of A:T transitions when cells grow on LB at 37°C may be due in part to suppression of transitions at G:C base pairs.

### BPS rates are dramatically influenced by the local sequence context

The large number of BPSs accumulated over the entire chromosome in the MMR defective strains allows us to analyze the local sequence context of the BPSs with great statistical power. To evaluate the sequence context of the BPSs we recorded the 19 bases 3′ and 5′ to each mutated base. These base sequences or their reverse complements were oriented so to flank the purine member of the mutated base pair. The rate at which each base appeared in each of the 38 flanking positions was calculated from all the experiments with MMR defective strains, normalized to the number of that base in the genome, and then divided by the overall sum of the bases. The resulting fraction of each base in each position is given in Supplementary Figure S1A and S1B, which reveals that at least 2 bases on each side of a base can influence its mutation rate. However, the immediately adjacent bases are most important, so our analysis considers only triplets with the target base in the center. While there are 64 possible triplets, in double-stranded DNA a triplet and its reverse complement (each read 5′ to 3′) are equivalent since each pairs with the other on the opposite DNA strand. As shown in Figure 2A and 2B, the presence of a C 3′ to the purine (= a G 5′ to the pyrimidine on the other strand) strongly influences the mutation rate. 5′N**A**C3′+5′G**T**N3′ triplets are hotspots for A:T mutations (throughout this report, triplets are displayed 5′ to 3′ with target base in the center and displayed in bold); the average mutation rate at these triplets was 12-fold higher than that of the other A:T-containing triplets. The mutation rate of A:T base pairs in most highly mutated triplet, 5′T**A**C3′+5′G**T**A3′, was 51 times greater than that of the A:T base pairs in the least mutated triplet, 5′A**A**A3′+5′T**T**T3′. 5′N**G**C3′+5′G**C**N3′ triplets are also minor hotspots for GC mutations; the mutation rate at these triplets was 2.4-fold higher than that at the other G:C containing triplets. As we previously reported from a smaller dataset (Sung *et al.* 2015), the G:C base pairs in the triplets 5′C**G**C3′+5′G**C**G3′ and 5′G**G**C3′+5′G**C**C3′ had the highest mutation rates. The mutation rate of G:C base pairs in the most highly mutated triplet, 5′G**G**C3′+5′G**C**C3′, was 17 times greater than that of the G:C base pairs in least mutated triplet, 5′A**G**T3′+5′A**C**T3′.

Ninety seven percent of the mutations at the 5′N**A**C3′+5 G**T**N3′ hotspots were transitions. So significant were A:T transitions at these triplets that they alone accounted for the 3-fold difference between transition rates at A:T and G:C base pairs in the MMR spectrum (Figure 1). Transitions at 5′N**A**C3′+5′G**T**N3′ triplets also accounted for most of the difference in A:T transition rates when the MMR defective strains were grown on LB versus minimal medium (Table 1). On minimal medium the average A:T transition rates at most triplets was reduced 2-fold, but at 5′N**A**C3′+5′G**T**N3′ triplets the rate was reduced 4-fold. However, these triplets were nonetheless hotspots; A:T transitions occurred at twice the rate at 5′N**A**C3′+5′G**T**N3′ triplets than at other triplets even when the MMR defective strains were grown on minimal medium (Supplemental Figure S2).

**Figure 2.**
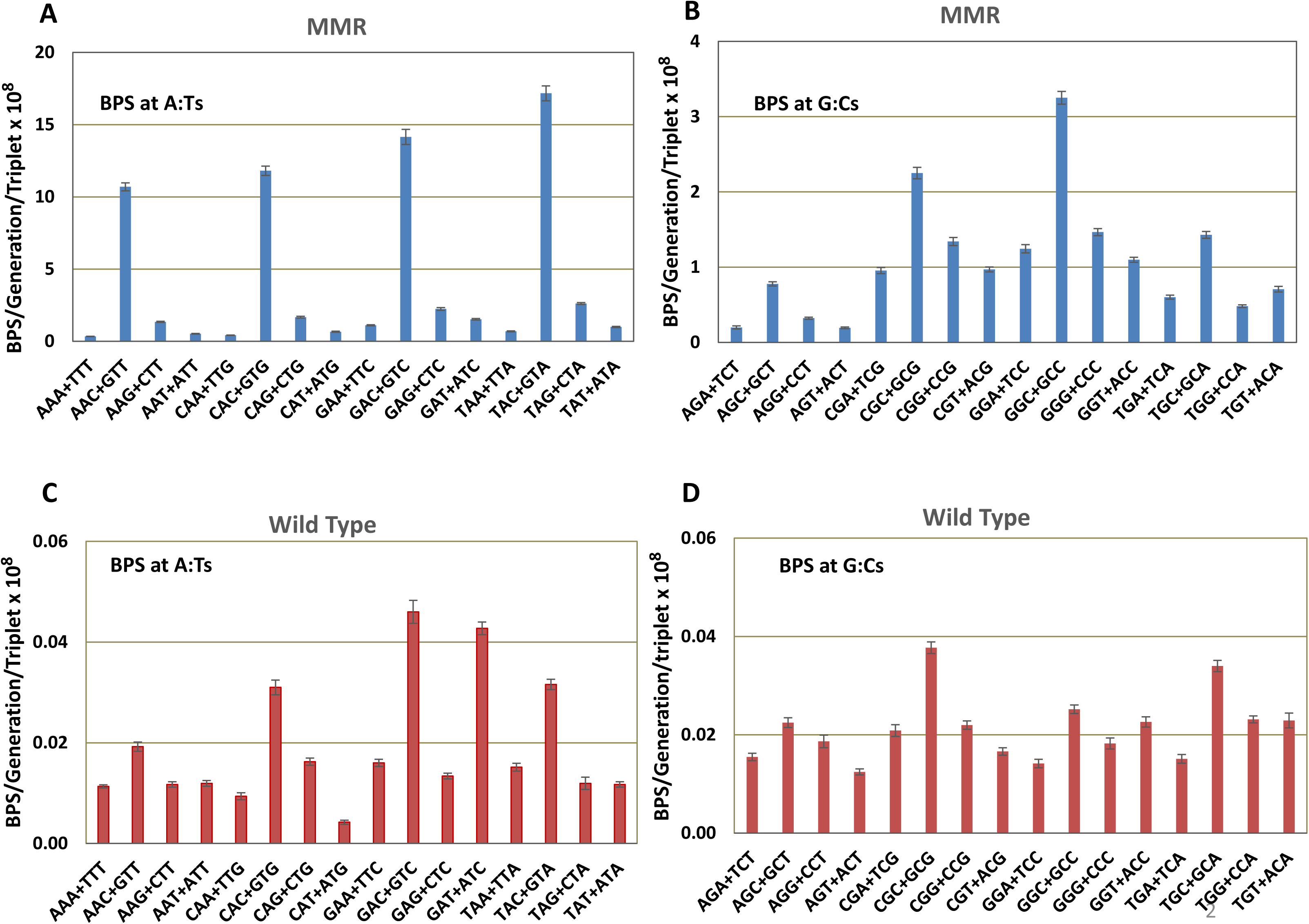
Base-pair substitution rates are influenced by the local sequence context. The data from the ten MA experiments with MMR-defective strains (A & B) and 8 MA experiments with MMR-competent strains (C & D) are combined to give the results labeled MMR and Wild Type (see Materials and Methods). The X-axis labels are the 32 sets of non-redundant triplets read 5′ to 3′ with the target base in the center of each triplet. Bars represent the mean BPS rates at each triplet; error bars are 95% CLs. Note the change in scale in the MMR charts between the BPS at A:Ts (A) and at G:Cs (B).

### Mononucleotide runs are hotspots for BPSs

The distribution of the 30,061 BPS over all 4,639,675 base pairs in the genome does not fit a Poisson distribution. With a mean of 0.00648 BPS/base pair, there should be only 97 base pairs mutated twice, and essentially none mutated more than twice. Instead, in the MMR-defective strains, independently-arising BPS occurred twice at 721 base pairs, three times at 30 base pairs, four times at 3 base pairs, and five times at 3 base pairs. To investigate this anomaly we examined the local sequence of all the base pairs that were mutated more than once. As shown in Supplemental Table 7, there was a strong bias for repeat mutations to be located at the ends of mononucleotide runs. Indeed, the base pairs mutated four and five times each were at the ends of runs ≥ 5 Nts. Strikingly, at two of the base pairs mutated 5 times there were 9 instances of the G:C to C:G transversion, a very rare mutation.

These results prompted us to evaluate BPSs associated with mononucleotide runs throughout the genome. If runs are not hotspots, the expected mutation rate per base pair is simply the average number of BPSs per base pair. However, as shown in Figure 3A, the rate of BPS associated with runs increased with the length of the run, especially at runs >6 Nt, reaching a maximum of 18.5-fold greater than expected at base pairs associated with runs of length 8 (there was only one mutation at a run 9 Nts, of which there are 19 times in the genome, and no mutations at the single run of 10 Nts in the genome). The majority of these BPS occurred adjacent to runs: of the 2040 BPSs associated with runs ≥4 Nts, 1280 (63%) occurred at the Nt next to the run whereas the rest occurred at the last base of the run.

**Figure 3.**
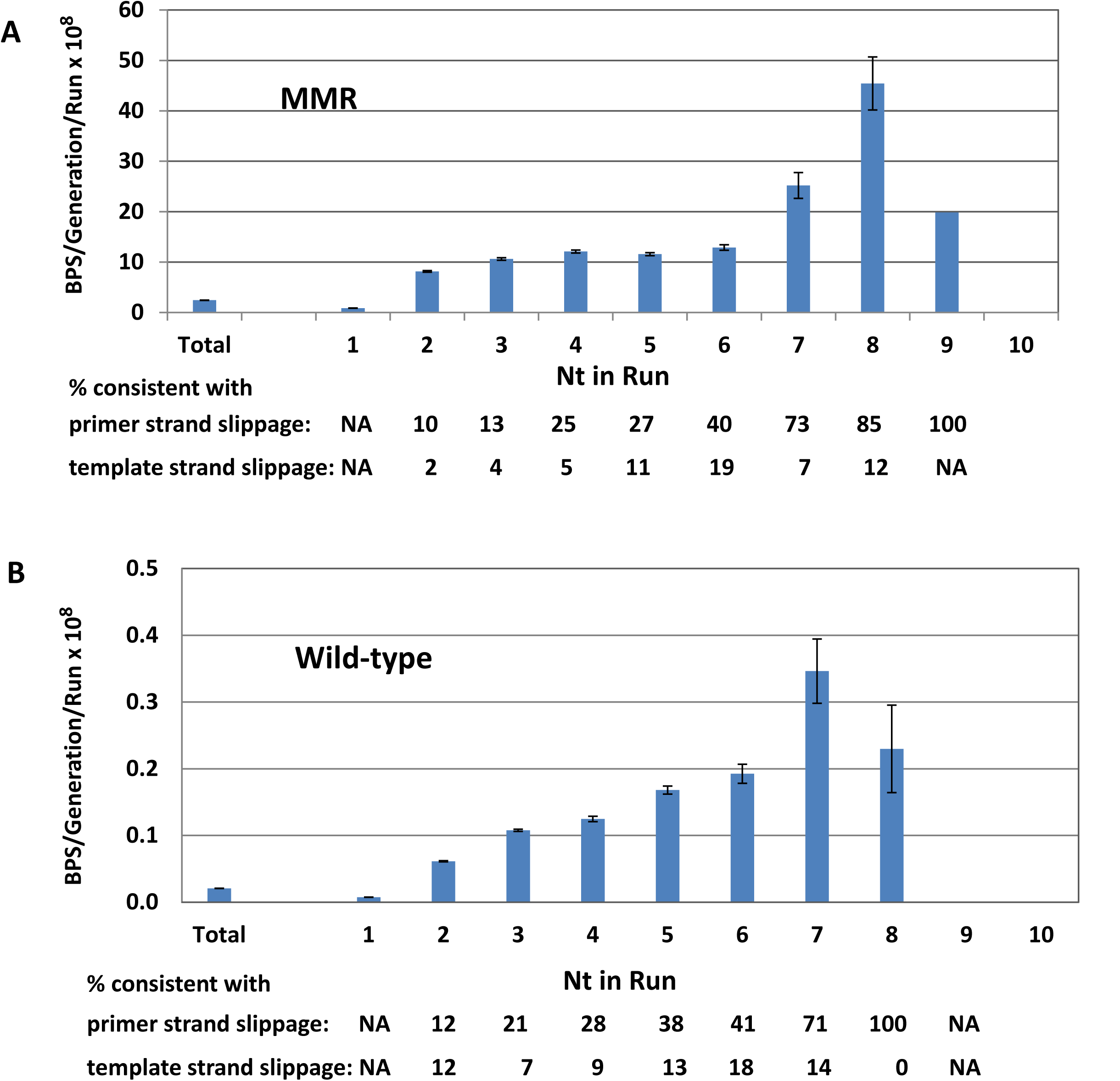
Mononucleotide runs are hotspots for BPSs in both MMR-defective (A) and MMR-proficient (B) strains. Bars represent the mean BPS rates per generation at basepairs adjacent to or within a mononucleotide run divided by the number of runs of each length in the genome. For BPS not associated with runs (*i.e*. Nt in run = 1) the BPS rate per generation was divided by the total number of Nt in the genome minus the Nts in runs. Total = all BPS/generation divided by all Nt in the genome. Fractions are the number of BPS that are consistent with either primer or template strand slippage divided by the total number of BPS that occurred associated with runs of each length. The data are from the ten MA experiments with MMR-defective strains (A) and 8 MA experiments with MMR-competent strains (B). Nt, nucleotide; NA, not applicable. Error bars are 95% CL.

BPSs associated with runs could occur by a mechanism known as transient misalignment or dislocation mutagenesis (Fresco and Alberts 1960; Fowler *et al.* 1974; Kunkel and Soni 1988). During DNA replication, either the template or the primer strand loops out but DNA polymerase continues replicating. When the strand realigns, it creates a mispair if the 3′ base of the primer does not match the 5’ base of the template. This mechanism is similar to that proposed for frameshift mutations (Streisinger *et al.* 1966) and is presumably enhanced at mononucleotide runs where DNA polymerases tend to slip.

By examining the nature of the mispairs, we tested whether this mechanism could explain the BPS associated with runs. If the primer strand slips out and realigns, it creates a BPS 5′ to the run that is templated by the run (Supplemental Figure S3A). Examination of the sequences showed that 585 (46%) of the 1280 BPSs that occurred adjacent to runs ≥4 Nts were consistent with this mechanism. If, instead, the template strand loops out and realigns, it creates a BPS at the last base of the run that is templated by the base 5′ to the run (Supplemental Figure S3B). However, only at runs ≥6 Nt was the number of BPSs consistent with this mechanism higher than the expected 33%. Of the 48 BPS that occurred at the last base of a run ≥6 Nt, 35 (73%) were consistent with being created by this mechanism. This analysis also revealed that the fraction of BPSs associated with runs that could be explained by transient misalignment increased with the length of the run.

However, it is clear from Figure 3A that not all of the BPSs associated with runs can be explained by transient misalignment. Also, of the cases of multiple mutations at a given base pair, in two cases, one each of three and five occurrences, one mutation of each set was different (specifically, GC to AT instead of GC to CG; Supplemental Table S7). In addition, 78% of the duplicate BPSs and 53% of the triplicate BPSs were at base pairs not associated with runs. Thus, other mechanisms must be responsible for many of these “super hotspots”. For example, the fidelity of polymerization and/or efficiency of proofreading may decline at runs.

### BPSs are DNA-strand biased

In a previous paper (Lee *et al.* 2012) we reported that in a strain deleted for *mutL*, A:T transitions were twice as frequent when A was on the lagging strand template (LGST) and T on the leading strand template (LDST) than when the mutated A:T was in the opposite orientation. In contrast, G:C transitions were twice as frequent when the C was on the LGST and G on the LDST than when the mutated G:C was in the opposite orientation. Similar results were obtained in MA experiments with a MMR-defective *Pseudomonas aeruginosa* strain (Dettman *et al.* 2016). The data presented here confirm and extend these findings. From the 10 experiments with MMR defective strains reported here (29,234 transitions recovered), the bias for A vs. T on the LGST was 2.44 ± 0.09, whereas the expected bias is 0.99 (χ^2^ = 2068, P ≈0); the bias for C vs. G on the LGST was 2.34 ± 0.09, whereas the expected bias is 0.94 (χ^2^ = 696, P ≈0) (ratios are means ± 95%CLs).

The strand bias in the MMR-defective strains varied with local sequence context (Figure 4A and 4B): among A:T transitions, the bias for A vs. T on the LGST varied 3-fold, with the highest ratio (3.17±0.13) at 5′A**A**C3′+5′G**T**T3′ sites, and the lowest (1.24±0.12) at 5′C**A**A3′+5′T**T**G3′ sites. Among G:C transitions, the bias for C vs. G on the LGST varied 5-fold, with the highest (3.30±0.19) at 5′G**G**T3′+5′A**C**C3′ sites and the lowest (0.65±0.08) at 5′A**G**G3′+5′C**C**T3′ sites (ratios are means ± 95%CLs).

**Figure 4.**
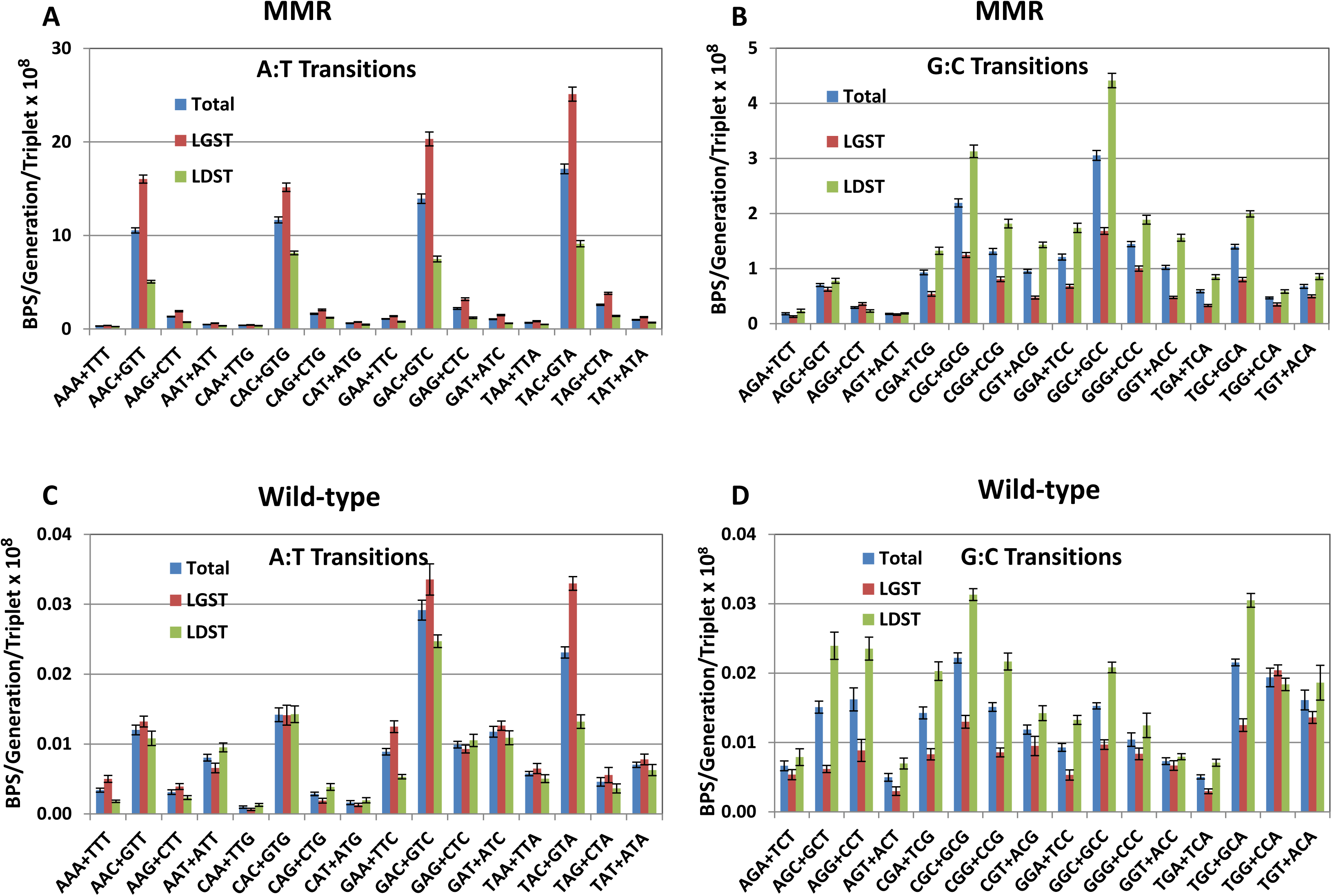
The DNA-strand bias of transition mutations varies strongly with sequence context in MMR-defective strains, but weakly in MMR-proficient strains. Bars represent the mean rates of transitions accumulated in 10 experiments with MMR-defective strains (A & B) and 8 experiments with MMR-proficient strains (C & D) (see Materials and Methods). Mutation rates per generation at each triplet were divided by the number of that triplet in the genome. Error bars are 95% CLs. The X-axis labels are the 32 sets of non-redundant triplets read 5′ to 3′ with the target base in the center of each triplet. Note the change in scale in the MMR charts between the transitions at A:Ts (A) and at G:C (B). LGST, the target purine as displayed was on the lagging strand template; LDST, the target purine as displayed was on the leading strand template.

In contrast to transitions, both A:T and G:C transversions in the MMR-defective strains were more likely to occur with the pyrimidine on the LGST and the purine on the LDST (Supplemental Figure S4). For A:T transversions the ratio was 1.68±0.13 instead of the expected 1.01 (χ^2^ = 17, P = 3×10^−5^); for G:C transversions the ratio was 1.15±0.11, but this ratio is not significantly different from expected 0.94 (χ^2^ = 1.5, P =0.22) (ratios are means ± 95%CLs). There were too few transversions to make meaningful conclusions about the influence of the local sequence on strand bias.

We previously speculated that the strand bias observed for A:T transitions was due to a greater tendency for DNA polymerase to insert a C opposite A when the A templates lagging-rather than leading-strand synthesis (Lee *et al.* 2012). To test this hypothesis we performed a MA experiment with a *mutL ndk* double mutant strain. The *ndk* gene encodes nucleoside diphosphate kinase, and, in its absence, the cellular concentration of dCTP increases at least 2-fold (Schaaper and\Mathews 2013) and as much as 12-fold (Maslowska *et al.* 2015). Relative to the MMR defective strains, loss of Ndk increased the BPS rate 5.4±0.1-fold overall, with the largest increases in A:T to G:C transitions (6.7±0.09-fold), and A:T to C:G transversions (25.6±0.6-fold) (values are means ± 95%CLs).

As expected, these enhanced BPS rates can be explained by an increase in the frequency at which Cs are inserted opposite template As and Ts. As shown in Table 2, in the *mutL ndk* double mutant strain, there was a ≈2-fold bias for the template A and T to be on the LGST (actual biases ± 95%CLs are 1.93± 0.003 for A and 1.57±0.003 for T; χ^2^ for the differences from the 0.99 and 1.01 expected biases are 640, P ≈ 0, and 13, P <0.001, respectively). This result supports the hypothesis that C is misinserted about twice as often during lagging-strand synthesis as during leading-strand synthesis. However, the same ratio would result if the proofreader removes the misinserted C twice as frequently when it is on the leading rather than the lagging strand.

**Table 2.**
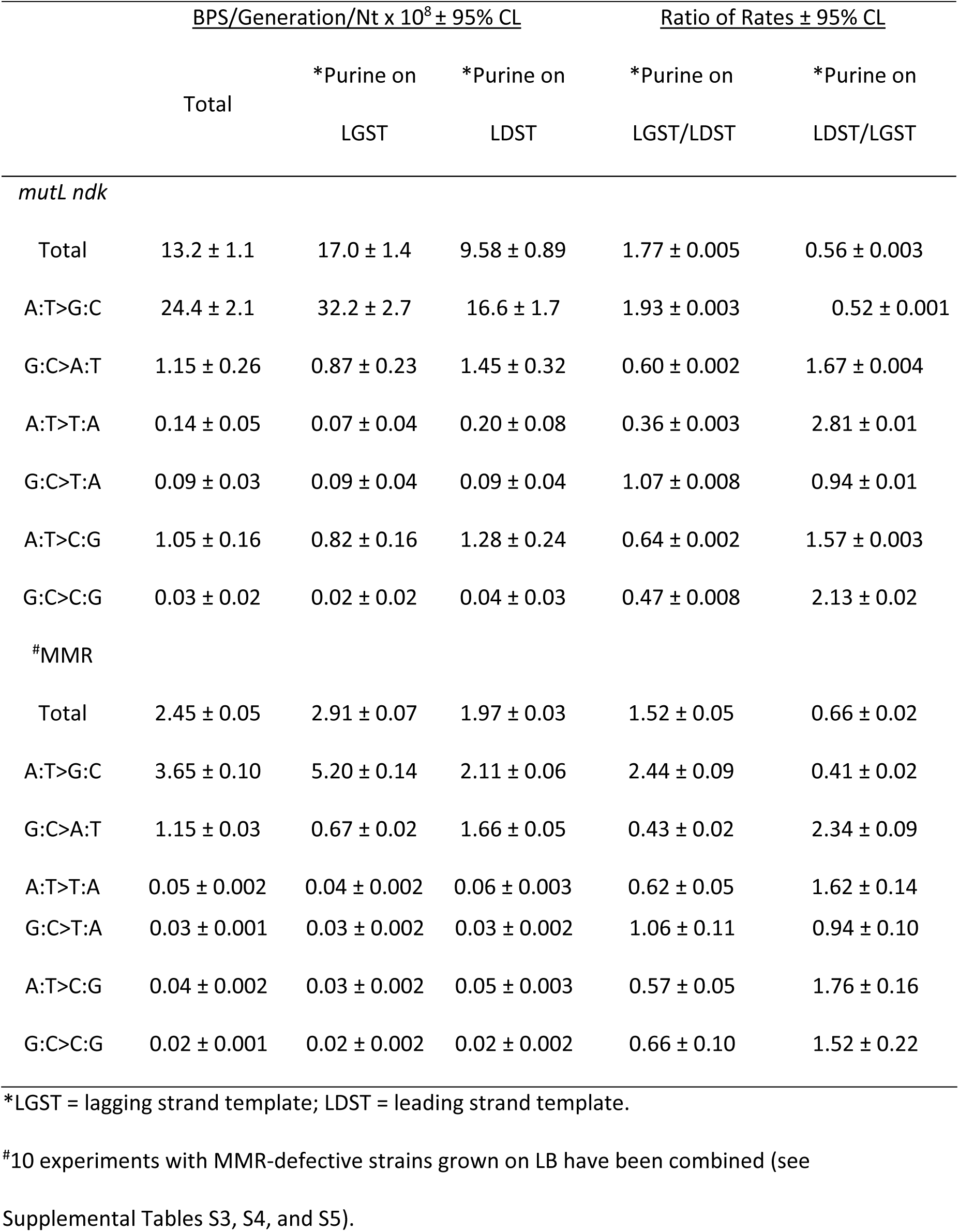
Rate and strand bias of the *mutL ndk* mutant strain compared to MMR-defective strains

|  |  | <u>BPS/Generation/Nt x 10<sup>8</sup> ± 95% CL</u> |  | <u>Ratio of Rates ± 95% CL</u> |  |
| --- | --- | --- | --- | --- | --- |
|  | Total | *Purine on<br>LGST | *Purine on<br>LDST | *Purine on<br>LGST/LDST | *Purine on<br>LDST/LGST |
| *mutL ndk* |  |  |  |  |  |
| Total | 13.2 ± 1.1 | 17.0 ± 1.4 | 9.58 ± 0.89 | 1.77 ± 0.005 | 0.56 ± 0.003 |
| A:T>G:C | 24.4 ± 2.1 | 32.2 ± 2.7 | 16.6 ± 1.7 | 1.93 ± 0.003 | 0.52 ± 0.001 |
| G:C>A:T | 1.15 ± 0.26 | 0.87 ± 0.23 | 1.45 ± 0.32 | 0.60 ± 0.002 | 1.67 ± 0.004 |
| A:T>T:A | 0.14 ± 0.05 | 0.07 ± 0.04 | 0.20 ± 0.08 | 0.36 ± 0.003 | 2.81 ± 0.01 |
| G:C>T:A | 0.09 ± 0.03 | 0.09 ± 0.04 | 0.09 ± 0.04 | 1.07 ± 0.008 | 0.94 ± 0.01 |
| A:T>C:G | 1.05 ± 0.16 | 0.82 ± 0.16 | 1.28 ± 0.24 | 0.64 ± 0.002 | 1.57 ± 0.003 |
| G:C>C:G | 0.03 ± 0.02 | 0.02 ± 0.02 | 0.04 ± 0.03 | 0.47 ± 0.008 | 2.13 ± 0.02 |
| #MMR |  |  |  |  |  |
| Total | 2.45 ± 0.05 | 2.91 ± 0.07 | 1.97 ± 0.03 | 1.52 ± 0.05 | 0.66 ± 0.02 |
| A:T>G:C | 3.65 ± 0.10 | 5.20 ± 0.14 | 2.11 ± 0.06 | 2.44 ± 0.09 | 0.41 ± 0.02 |
| G:C>A:T | 1.15 ± 0.03 | 0.67 ± 0.02 | 1.66 ± 0.05 | 0.43 ± 0.02 | 2.34 ± 0.09 |
| A:T>T:A | 0.05 ± 0.002 | 0.04 ± 0.002 | 0.06 ± 0.003 | 0.62 ± 0.05 | 1.62 ± 0.14 |
| G:C>T:A | 0.03 ± 0.001 | 0.03 ± 0.002 | 0.03 ± 0.002 | 1.06 ± 0.11 | 0.94 ± 0.10 |
| A:T>C:G | 0.04 ± 0.002 | 0.03 ± 0.002 | 0.05 ± 0.003 | 0.57 ± 0.05 | 1.76 ± 0.16 |
| G:C>C:G | 0.02 ± 0.001 | 0.02 ± 0.002 | 0.02 ± 0.002 | 0.66 ± 0.10 | 1.52 ± 0.22 |

\*LGST = lagging strand template; LDST = leading strand template.

In the *mutL ndk* mutant strain the rate of A:T > G:C transitions, presumably resulting from C inserted opposite template A, was about 25-fold greater at 5′N**A**C3′ triplets than at the other A containing triplets (Supplemental Figure S5). Interestingly, these same triplets were hotspots for A:T > C:G mutations. Assuming these transversions occurred because C was inserted opposite template T in the complementary 5′N**T**G 3′ triplet, the misinsertion occurred during leading strand replication. In this orientation, the excess of dCTP in the *ndk mutL* strain would strongly promote the formation of the G:C basepair 3’ to the T:C mispair, which would tend to protect the mispair from editing by proofreader. This “next nucleotide” effect is a well-known property of proofreading DNA polymerases (Fersht 1979; Bloom *et al.* 1997). Thus, the favorable orientation for the mispair may overcome the normal fidelity of leading strand synthesis.

### Alternative Explanations for the BPS spectrum and strand bias

It is possible that some of our results could be due to the appearance in the DNA of base analogues with alternative base-pairing properties. For example, deoxyinosine triphosphate (dITP) is produced by deamination of dATP and, if incorporated into the DNA, will basepair with dTTP and, more stably, dCTP (ALSETH *et al.* 2014). Thus, if incorporated opposite template T, dITP can cause A:T to G:C transitions if it subsequently pairs with C. Deamination of adenine in the DNA produces hypoxanthine, which preferentially pairs with cytosine, also causing A:T to G:C transitions. dITP is cleared from the nucleotide pool by the RdgB protein (Chung *et al.* 2002), and hypoxanthine is excised from DNA by a pathway initiated by endonuclease V (Nfi) (CAO 2013). Previous studies have shown that neither the loss of Nfi nor of RdgB has a significant mutagenic consequence, suggesting that these base analogues do not contribute to spontaneous mutations (Budke and Kuzminov 2006; Foster *et al.* 2015). To test if these base analogues are important in the MA experiments reported here, we used fluctuation tests to compare the mutation rates of Δ*rdgB* Δ*mutL* and Δ*nfi* Δ*mutL* mutant strains to that of the Δ*mutL* parental strain. Loss of neither enzyme had a mutagenic effect (Supplemental Table S8).

DNA polymerase I (Pol I) is responsible for maturation of the Okasaki fragments produced during lagging strand DNA synthesis. Thus, the bias for A:T transitions to occur when A is on the LGST could be due to Pol I inserting C’s opposite As during this process or, possibly, during repair synthesis if Pol III disassociates. However, this hypothesis seems unlikely since a Pol I mutant lacking its proofreading function, which has at least a 7-fold greater error-frequency than the wild-type Pol I (Bebenek *et al.* 1990), had only a modest effect on spontaneous mutation rates (Makiela-Dzbenska *et al.* 2009; Makiela-Dzbenska *et al.* 2011). These authors concluded that Pol I’s only role in replication is to accurately fill in the gaps left by Okasaki fragment maturation (Fijalkowska *et al.* 2012).

### Mutational features found in MMR-defective strains are also evident in wild-type strains

The low BPS rate of wild-type strains (≈120-fold lower than that of MMR-defective strains) makes it difficult to accumulate sufficient numbers of mutations to allow as detailed analysis of mutational features as presented above. However, by combining the results obtained for the wild-type strain (Lee *et al.* 2012) with those obtained from MA experiments performed with strains carrying DNA repair defects that did not affect the mutational rate or spectrum (Foster *et al.* 2015), we were able to achieve 1,933 BPSs from eight independent experiments for analyses (see Materials and Methods).

The strong sequence context biases seen in the spectrum from the MMR-defective strains are also evident in the wild-type spectrum (Figure 2C and 2D). In particular, 5′N**A**C3′+5′G**T**N3′ triplets are hotspots for A:T mutations, particularly transitions. 5′G**A**T3′+5′A**T**C3′ triplets are also hotspots, but these are entirely due to transversion mutations occurring at G**A**TC sequences. *E. coli*’s Dam methylase methylates the 6-position of adenines in GATC sequences, making the adenine prone to depurination, which results in transversion mutations (Lee *et al.* 2012). These sites are also hotspots for transversions in the MMR-defective strains, but the dominance of transition mutations obscures this result (Figure 2A). In fact, in the MMR-defective strains the rate of A:T transversions at G**A**TC sites was 8-fold higher than at all other A:T base pairs, but in the wild-type strains it was only 4-fold higher, meaning that MMR prevents many mutations at these sites.

Also, as in the MMR-defective strains, in the wild-type strains BPSs associated with mononucleotide runs occurred at rates far greater than expected (Figure 3B). The rate of BPSs increased with the length of the run, reaching a maximum of 17.5-fold greater than expected at nucleotides associated with runs of length 7 (in the wild-type strains there was only 1 mutation at runs of 8 Nt, which occur 216 times in the genome, and no mutations at longer runs). Of the 179 BPS associated with runs ≥4 Nts, 81 (45%) were consistent with being due to transient misalignment.

The DNA-strand bias of transitions mutations accumulated in the wild-type strains was not as strong as the strand bias in the MMR-defective stains. From the 8 experiments with MMR proficient strains (1052 transitions recovered), the bias for C vs. G on the LGST was 1.91 ± 0.05, which is significantly greater than the expected 0.94 (χ^2^ = 39, P ≈ 0). However, the bias for A vs. T on the LGST was 1.23 ± 0.05, which is not significantly different from the 0.99 expected (χ^2^ = 2, P = 0.12) (ratios are means ± 95%CLs). Although the biases among A:T transitions varied with sequence context (Figure 4C and 4D), only at 5′T**A**C3′+5′G**T**A3′ sites, where the ratio of A vs. T on the LGST was 2.5 ± 0.2, were enough mutations recovered so that the bias was significantly greater than expected (χ^2^ = 4, P = 0.03) (ratios are means ± 95%CLs). A similar reduction of strand bias when MMR was active was also seen in MA experiments with a *Pseudomonas aeruginosa* strain (Dettman *et al.* 2016).

### The efficiency of MMR is influenced by the local sequence context

One explanation for the sequence bias of BPS rates is that the efficiency of MMR is low in certain sequence contexts, e.g. 5′G**A**C3′+5′G**T**C3′ triplets, and thus mutation rates are high at those sites in wild-type strains. However, this is not the case. MMR efficiency can be estimated by dividing the mutation rate at a given sequence context determined in MMR defective strains by that determined in MMR proficient strains. As shown in Figure 5, when this is done the efficiency of MMR varies 43-fold with sequence context. Importantly, the 5′N**A**C3′+5′G**T**N3′ hotspots are among the most efficiently repaired by MMR; thus, the fact that these sites are hotspots in the wild-type strains is not because they are poor substrates for MMR. In contrast, mismatches associated with mononucleotide runs are not particularly good substrates for MMR. MMR efficiency does not depend on run length and the MMR/WT ratio for BPSs associated with runs ≥4 Nts was 87 ± 3 (mean ± 95% CL), somewhat less than the 120-fold ratio calculated for all BPSs.

**Figure 5.**
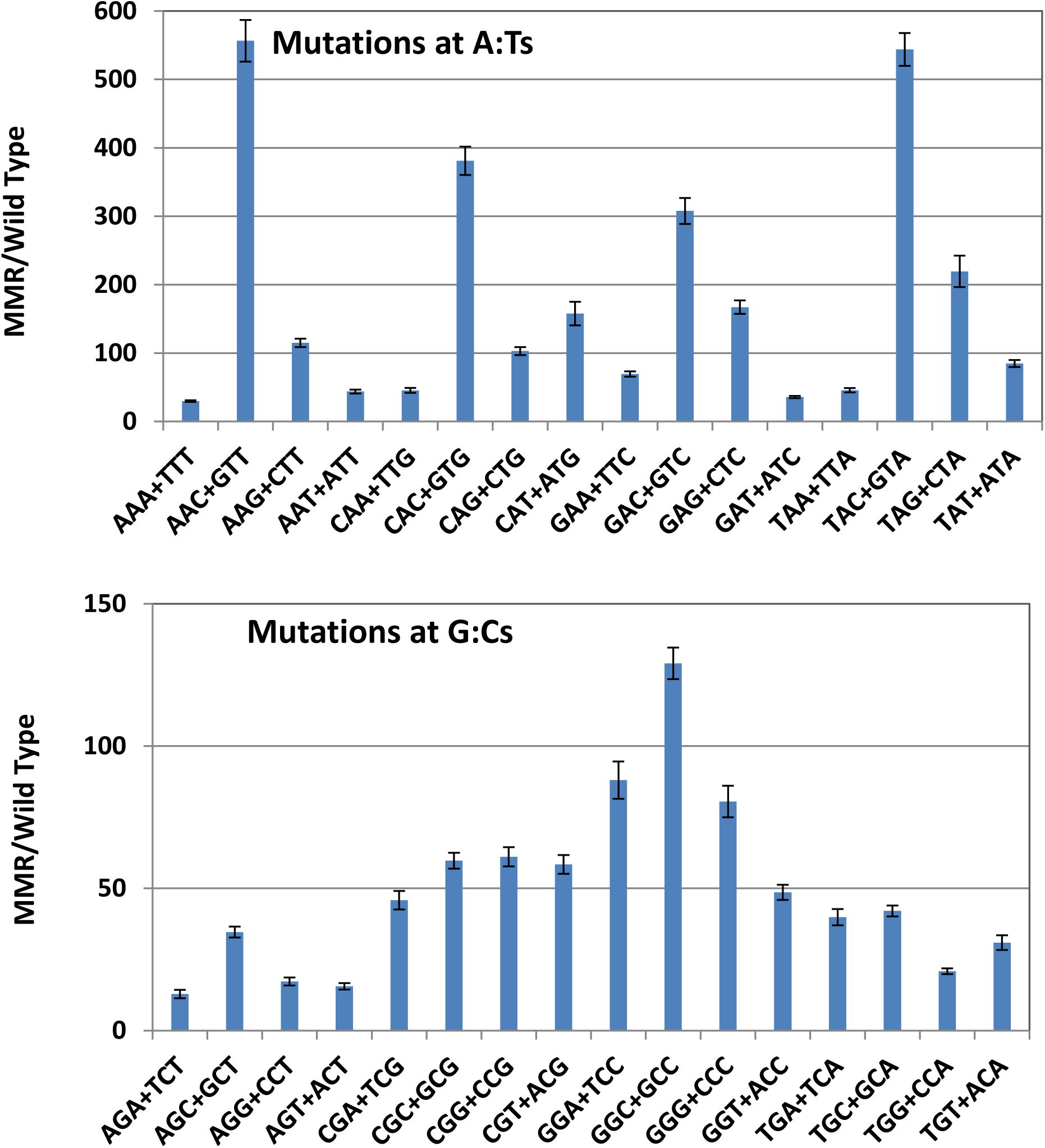
The efficiency of MMR is influenced by the local sequence context. Bars represent the mean ratio of mutation rates at each triplet in MMR defective versus MMR proficient strains. Error bars are 95%CL calculated for the ratio. The X-axis labels are the 32 sets of non-redundant triplets read 5′ to 3′ with the target base in the center of each triplet. Note the change in scale between the results of mutations at A:Ts and at G:C.

## DISCUSSION

The observations presented here can be summarized as follows:

1. In MMR-defective strains, the spectrum of BPSs is dominated by A:T transitions, which occur at a rate 3-fold greater than the rate of G:C transitions, and 75 to nearly 200-fold greater than the rates of all other BPSs. The preponderance of A:T transitions over G:C transitions is almost entirely due to mutations at 5′N**A**C3′+5′G**T**N3′ sites.
2. The overall rate of BPSs in MMR-defective strains is about 120-fold greater than in MMR-proficient strains. At A:T sites the increase is nearly 200-fold, whereas at G:C sites it averages only 50-fold.
3. The increase in BPS rate with the loss of MMR varies with local sequence context. Among A:T sites, the mutation rates at 5′N**A**C3′+5′G**T**N3′ sites are increased 300-500 fold, whereas the mutation rates at 5′N**A**A3′+5′T**T**N3′ sites are increased only 30-70 fold. Among G:C sites, the variation increases of the mutation rate is smaller, but, nonetheless, the range is 10-fold, with a maximum of 130-fold at 5′G**G**C3′+5′G**C**C3′ sites and a minimum of 13-fold at 5′A**G**A3′+5′T**C**T3′ sites.
4. When MMR-defective strains are grown on minimal medium, the BPS rate is reduced 2-fold and the rates of G:C and A:T transitions are equalized. Slowing the growth rate of the cells by incubating them on LB medium at low temperature reduces the overall BPS rate by 60% but has a 2-fold greater effect on A:T than on G:C transitions. Thus, growth rate appears to be an important factor in the production of A:T transitions, but LB also may also promote A:T and suppress G:C transitions.
5. Base-pairs adjacent to or within mononucleotide runs are particularly vulnerable to mutation in both MMR-defective and MMR-proficient strains. Many, but not all, of these BPS appear to be due to transient misalignment of either the primer or the template DNA strand during replication.
6. In MMR-defective strains there is a two-fold bias for A:T transitions to occur when the A is on the lagging strand template (LGST) and a two-fold bias for G:C transitions to occur when the C is on the LGST. Only the G:C bias is significant in wild-type strains. Increasing the cellular concentration of dCTP by deleting the *ndk* gene resulted in an increase in BPSs consistent with misinsertion of C opposite template As and Ts. These occurred about two-fold more frequently when the A or T was on the LGST rather than the LDST.
7. Over all BPSs, the efficiency of MMR correction varies 43-fold with sequence context. The mismatches leading to the high rate of transitions at 5′N**A**C3′+5′G**T**N3′ sites are corrected by MMR 9-fold more efficiently than the mismatches leading to the low rate of transitions at 5′N**G**A/T3′+5′T/A**C**N3′ sites.

That MMR correction is influenced by both the nature of the mismatch and its sequence context has long been appreciated (Cox *et al.* 1972; Modrich 1991). Thus, it is not surprising that previous studies of the mutational consequences of loss of MMR, which have used a variety of mutational targets, have produced some divergent results. However, all previous reports have identified transitions as the major BPS enhanced by the loss of MMR. By using an unbiased protocol with the entire genome as the mutation target, we have confirmed these previous results and extended them with statistical power hitherto not possible.

Our finding that 5′N**A**C3′+5′G**T**N3′ sites are hotspots for transitions could not have been seen in studies using reversion assays because such assays are limited to one, or a few, mutational targets. But our results were presaged by studies that employed mutational targets that, while still limited to a single gene, were broad enough to reveal the influence of local sequence context. Using mutation of the *lacI* gene to give the LacI^−d^ phenotype, Schaaper and Dunn, 1987, found that A:T transitions were 2-fold more frequent than G:C transitions in MMR-defective strains, and 88% of the A:T transitions occurred at seven 5′N**A**C3′+5′G**T**N3′ sites (Schaaper and Dunn 1987). Likewise, using knockout mutations of the bacteriophage P22 *mnt* gene, Wu *et al.*, 1990, found that A:T transitions were nearly 4-fold more frequent than G:C transitions in MMR-defective strains; two 5′N**A**C3′+5′G**T**N3′ triplets accounted for 36% of the A:T transitions, although one 5′G**A**G3′+5′C**T**C3′ site was equally mutated (Wu *et al.* 1990). Finally, using mutation to Rif^R^, which occurs by BPSs at 80 sites in the *rpoB* gene, Garibyan *et al.*, 2003, found that 56% of the A:T transitions in a *mutS* mutant strain occurred at one 5′G**A**C3′+5′G**T**C3′ triplet (Garibyan *et al.* 2003). However, in all of these studies not every 5′N**A**C3′+5′G**T**N3′ triplet that could be mutated was mutated. In addition, among the seven 5′N**A**C3′+5′G**T**N3′ sites at which AT transitions were recorded in the *lacI−^d^* data, there was an 8-fold difference in mutation frequency (Schaaper and Dunn 1987). From our results, of the 512,268 5′N**A**C3′+5′G**T**N3′ triplets in the genome, 3% were mutated once and 0.1% were mutated more than once. Even disregarding the BPSs associated with mononucleotide runs, twice as many 5′N**A**C3′+5′G**T**N3′ sites than expected were mutated more than once. Thus, even within a mutation-prone sequence context, features in addition to the adjacent bases must influence the mutation rate of a given base pair. However, analysis of 19 bases on each side of each BPS did not reveal any additional sequence contexts influencing the probability of a given base being repeatedly mutated (Supplemental Figure S6)

The influences of local sequence context observed in our MA experiments have similarities and differences with the results of MA experiments with other bacteria. Mutations at 5′N**A**C3′+5′G**T**N3′ triplets dominated the BPSs in MA experiments with a MMR-defective *Pseudomonas aeruginosa* strain (Dettman *et al.* 2016), but in MA experiments with *Bacillus subtilis*, both Sung *et al.*, 2015, (Sung *et al.* 2015) and Schroeder *et al.*, 2016, (Schroeder *et al.* 2016) found that the most highly mutated triplets in both MMR-defective and wild-type strains were 5′C**A**C3′+5′G**T**G3′, 5′C**G**C3′+5′G**C**G3′, and 5′C**G**G3′+5′G**C**C3′. While 5′C**A**C3′+5′G**T**G3′ and 5′C**G**C3′+5′G**C**G3′ (but not 5′C**G**G3′+5′G**C**C3′) were frequently mutated in the experiments with *E. coli* reported here, A:T transitions at all 5′N**A**C3′+5′G**T**N3′ sites dominated the sequence context spectrum in both wild-type and, particularly, MMR-defective strains (Figure 2). But it is clear from these comparisons that G:C base pairs adjacent to the target base increase the probability of its mutation. Indeed, long ago the triplet 5′G**G**C3′+5′G**C**C3′ was identified as a transition hotspot in the *rII* gene of bacteriophage T4 (Singer 1984). G:C basepairs surrounding a base could potentiate misinsertion by DNA polymerase through base-stacking interactions (Kool 2001), and/or, because of their strong base-pairing, could prevent removal of the mismatch by inhibiting dissociation and transfer of the primer strand to the active site of the 3’exonuclease (Petruska and Goodman 1985; Bloom *et al.* 1994). Lowering the temperature and, thus, slowing down polymerization, would tend to counter these effects by giving proofreader more time to compete with the forward synthesis by polymerase, as has been observed when polymerization is slowed by lowering dNTP levels (Clayton *et al.* 1979). However, 5′N**A**C3′+5′G**T**N3′ sites are also hotspots in strains defective for proofreading, which suggests that polymerase has an intrinsic propensity for making errors at these sites (see Niccum *et al*, accompanying paper). A slower growth rate could counter this tendency by allowing polymerase more time to correctly discriminate between correct and incorrect incoming nucleotide.

Mononucleotide runs are hotspots for frameshift mutations (Streisinger *et al.* 1966; Kunkel 1986), and we previously reported that in our MA experiments the indel rate increases exponentially with the length of the run (Lee *et al.* 2012). The majority of such mutations can be attributed to polymerase slippage, *i.e.* during replication DNA polymerase allows either the template or the nascent (primer) strand to loop out and then continues synthesis, producing either an insertion or a deletion of a base. Our data show that mononucleotide runs are also hotspots for BPSs – in both MMR-proficient and MMR-deficient strains base pairs adjacent to runs ≥4 Nt long were 2- to 8-fold more likely to be mutated than expected and, in general, the mutation rate increased with run length (Figure 3). About half of these mutations could have been due to transient misalignment (Fresco and Alberts 1960; Fowler *et al.* 1974; Kunkel and Soni 1988), *i.e*. slippage of the primer or template strand at the repeat followed by insertion of a base and realignment, placing the incorrect base opposite the template base adjacent to the run, in the case of primer loop-out, or at the end of the run, in the case of template loop-out. Replication would then continue, preserving the mismatch, which, if uncorrected by proofreading, would become a mutation in the next round of replication (Supplemental Figure S3). However, the mechanism producing BPSs at the other half of the hotspots remains a mystery. Possibly replicating the “slippery” DNA affects the polymerase fidelity at adjacent bases.

The transitions that occur in MMR-defective strains are strand biased. We previously reported that the 2-fold bias for G:C transitions to occur when C is on the LGST rather than the LDST was likely due to cytosines deaminating to uracils, which code as thymines, on the single-stranded LGST during replication (Bhagwat *et al.* 2016). However, we have no such explanation for the 2-fold bias for A:T transitions to occur when A is on the LGST, although we speculated that DNA polymerase was more likely to insert a C opposite A when the A was templating lagging-strand synthesis (Lee *et al.* 2012). In contrast, other studies have indicated that leading strand DNA synthesis is more error-prone than lagging strand synthesis (Gawel *et al.* 2014). The results presented here with a strain defective for both MMR and Ndk support our original hypothesis, suggesting that lagging-strand synthesis is more error prone, at least when C is misinserted opposite template A or T. However, the mutational spectrum seen in the absence of MMR is the result of replication errors that are not corrected by the proofreader. Thus the strand bias we observe could be a result of the proofreader preferentially removing a misinserted nucleotide when it is on the leading strand, a hypothesis we explore in the accompanying paper.

## ACKNOWLEDGEMENTS

We thank the following past members of the P.L.F. laboratory for technical assistance: H. Bedwell-Ivers, C. Coplen, M. Durham, J. Eagan, J. Ferlmann, N. Gruenhagen, T. Gruenhagen, J. Healy, N. Ivers, C. Klineman, R. Meyer, R. Newlon, D. Osiedki, S. Patel, I. Rameses, L. Rich, S. Riffert, H. Rivera, D. Simon, K. Smith, B. Souders, K. Storvik, L. Tran, L. Whitson, B. Wojcik, N. Yahaya, and A. Ying Yi Tan. We also thank the anonymous reviewers of this manuscript for suggestions. The National BioResource Project at the (Japanese) National Institute of Genetics provided bacterial strains and plasmids. This research was supported by US Army Research Office Multidisciplinary University Research Initiative (MURI) Award [W911NF-09-1-0444 to P.L.F. & H.T.] and the National Institutes of Health [T32 GM007757 to B.A.N].

